# Fusogenicity and neutralization sensitivity of the SARS-CoV-2 Delta sublineage AY.4.2

**DOI:** 10.1101/2022.01.07.475248

**Authors:** Nell Saunders, Delphine Planas, William Bolland, Christophe Rodriguez, Slim Fourati, Julian Buchrieser, Cyril Planchais, Matthieu Prot, Isabelle Staropoli, Florence Guivel-Benhassine, Françoise Porrot, David Veyer, Hélène Péré, Nicolas Robillard, Madelina Saliba, Artem Baidaliuk, Aymeric Seve, Laurent Hocqueloux, Thierry Prazuck, Hugo Mouquet, Etienne Simon-Lorière, Timothée Bruel, Jean-Michel Pawlotsky, Olivier Schwartz

## Abstract

SARS-CoV-2 lineages are continuously evolving. As of December 2021, the AY.4.2 Delta sub-lineage represented 20 % of sequenced strains in UK and has been detected in dozens of countries. It has since then been supplanted by the Omicron variant. AY.4.2 displays three additional mutations (T95I, Y145H and A222V) in the N-terminal domain (NTD) of the spike when compared to the original Delta variant (B.1.617.2) and remains poorly characterized. Here, we analyzed the fusogenicity of the AY.4.2 spike and the sensitivity of an authentic AY.4.2 isolate to neutralizing antibodies. The AY.4.2 spike exhibited similar fusogenicity and binding to ACE2 than Delta. The sensitivity of infectious AY.4.2 to a panel of monoclonal neutralizing antibodies was similar to Delta, except for the anti-RBD Imdevimab, which showed incomplete neutralization. Sensitivity of AY.4.2 to sera from individuals having received two or three doses of Pfizer or two doses of AstraZeneca vaccines was reduced by 1.7 to 2.1 fold, when compared to Delta. Our results suggest that mutations in the NTD remotely impair the efficacy of anti-RBD antibodies. The temporary spread of AY.4.2 was not associated with major changes in spike function but rather to a partially reduced neutralization sensitivity.

## Introduction

The pandemic circulation of SARS-CoV-2 is associated with emergence of variants with increased inter-individual transmission or immune evasion properties. The Delta Variant of Concern (VOC), originally identified in India in 2020, has supplanted pre-existing strains worldwide in less than 6 months ^1 2^. The spike protein of Delta contains 9 mutations, when compared to the B.1 ancestral strain (D614G), including five changes in the NTD (T19R, G142D, Δ156, Δ157, R158G), two in the receptor binding domain (RBD) (L452R, T478K), one mutation close to the furin cleavage site (P681R) and one in the S2 region (D950N) ^3^. This set of mutations reduces sensitivity to antibody neutralization, enhances the fusogenicity of the spike and improves viral fitness ^3 4 5 6 7 8^. The increased transmissibility of VOCs may also be due to mutations in other viral proteins, such as R203N in the nucleocapsid (N) ^9^.

The Delta lineage is heterogeneous and continues to evolve. It can be divided into sublineages or clades ^10 11 12^. Different classifications exist. Next strain has classified the Delta variant into 3 main clades (21A, 21I and 21J). The Pangolin nomenclature is more resolutive and has designed almost 180 sublineages within these clades, all named AY as aliases to the B.1.617.2 lineages ^13^. Mutations fixed in one sublineage (e.g. spike: T19R, G142D or D950N) are also present at low frequencies in other sublineages.

This may reflect founder effects or similar selective pressures on these variants. One sublineage, termed AY.4.2 (or VUI-21OCT-01) has drawn attention due to its slow but continuous rise in UK between July and December 2021 ^14 15^. AY.4.2 sequences from 45 countries have been uploaded to the GISAID database. As of Dec 18, 2021, about 62,000 genomes have been reported in the UK on GISAID, representing about 15% of reported Delta cases in this country between December 1 and 18, 2021. Its occurrence has since then strongly diminished, as the Delta lineages have been replaced by Omicron strains worldwide ^16,17 18^.

The AY.4.2 sub-lineage is notably defined by the presence of Y145H and A222V mutations that lie within the NTD. Their impact on spike function is poorly characterized. Through modelling, the Y145H substitution has been predicted to decrease spike stability, but this has not been experimentally demonstrated ^19^. The mutation is located in close proximity to residue 144, which is deleted in the Alpha variant. A 141-144 deletion has also been reported in several chronically SARS-CoV-2 infected immunocompromised individuals ^20^. Furthermore, a 143-145 deletion is also observed in the Omicron variant ^21^. Deletions of aa 144 and adjacent residues may drive antibody escape ^22,23^. The A222V mutation was noted in the B.1.177 (or 20A.EU1) lineage that emerged in Spain and spread throughout Europe in summer 2020 ^24^. This lineage did not have obvious transmission advantage and its spread was mostly explained by epidemiological factors such as travelling ^24^. When introduced into the D614G spike, the A222V substitution slightly but not significantly impacted neutralization of pseudoviruses by human convalescent sera ^25^. The effect of combined Y145H and A222V mutations on the Delta background remains unknown. Of note, most AY.4.2 sequences (93%) now include the T95I mutation in the NTD of the spike, a substitution that was rarely observed in the original Delta B1.617.2 lineage, but which gradually appeared and is now present in 40% of Delta sequences on GISAID. The T95I substitution was previously detected in the close B.1.617.1 lineage (also termed Kappa) ^26^. It was also present in the B.1.526 lineage (also termed Iota) that accounted for up to 30% of sequenced cases in New York City in early 2021 ^27^. It is also present in the Omicron variant ^21^. This substitution was found in two vaccinated individuals with breakthrough infection and selected in an immunocompromised individuals with chronic COVID-19 treated with convalescent plasma and monoclonal antibodies ^28 29^. The T95 residue is located outside the NTD antigenic supersite and its contribution to immune evasion is poorly characterized ^26^.

Here, we studied the AY.4.2 spike by assessing its fusogenic activity, affinity to ACE2 and recognition by antibodies. We also isolated an infectious AY.4.2 strain and examined its sensitivity to a panel of monoclonal antibodies and sera from individuals having received two or three vaccine doses.

## Methods

No statistical methods were used to predetermine sample size. The experiments were not randomized and the investigators were not blinded to allocation during experiments and outcome assessment. Our research complies with all relevant ethical regulation.

### Orléans Cohort of convalescent and vaccinated individuals

Since August 27, 2020, a prospective, monocentric, longitudinal, interventional cohort clinical study enrolling 170 SARS-CoV-2-infected individuals with different disease severities, and 30 non-infected healthy controls is on-going, aiming to describe the persistence of specific and neutralizing antibodies over a 24-months period. This study was approved by the ILE DE FRANCE IV ethical committee. At enrolment, written informed consent was collected and participants completed a questionnaire which covered sociodemographic characteristics, virological findings (SARS-CoV-2 RT-PCR results, including date of testing), clinical data (date of symptom onset, type of symptoms, hospitalization), and data related to anti-SARS-CoV-2 vaccination if ever (brand product, date of first and second vaccination). Serological status of participants was assessed every 3 months. Those who underwent anti-SARS-CoV-2 vaccination had weekly blood and nasal sampling after first dose of vaccine for a 2 months period (ClinicalTrials.gov Identifier: NCT04750720). For the present study, we selected 56 convalescent and 28 vaccinated participants (16 with Pfizer and 12 with AstraZeneca). Study participants did not receive any compensation.

### Plasmids

A codon-optimized version of the reference Wuhan SARS-CoV-2 Spike (GenBank: QHD43416.1) was ordered as a synthetic gene (GeneArt, Thermo Fisher Scientific) and was cloned into a phCMV backbone (GeneBank: AJ318514), by replacing the VSV-G gene. The mutations for Alpha and Delta were added in silico to the codon-optimized Wuhan strain and ordered as synthetic genes (GeneArt, Thermo Fisher Scientific) and cloned into the same backbone. The D614G spike plasmid was generated by introducing the mutation into the Wuhan reference strain via Q5 site-directed mutagenesis (NEB). The T95I, Y145H and A222V were successively introduced into the Delta spike by the same process. Plasmids were sequenced prior to use. The primers used for sequencing and the site-directed mutagenesis are listed in the tables S3A and S3B.

### Cell-cell fusion assay

For cell–cell fusion assays, 3.5*10^5^ 293T cell lines stably expressing GFP1-10 were transfected in suspension with 50 ng of phCMV-SARS-CoV2-spike and 450 ng of pQCXIP-Empty for 30 min at 37°C. Cells were washed twice. For imaging, they were seeded at a confluency of 3.10^4^ cells per well in a 96 well plate. Vero GFP-11 cells were added at a confluency of 1.5.10^4^ cells per well.

The GFP area and the number of nuclei were quantified 18h post-transfection using Harmony High-Content Imaging and Analysis Software, as previously described ^37,38^. For surface staining they were seeded at a confluency of 6*10^4^ cells per well, and stained as described below using mAb 129.

### S-Fuse neutralization assay

U2OS-ACE2 GFP1-10 or GFP 11 cells, also termed S-Fuse cells, become GFP+ when they are productively infected by SARS-CoV-2 ^37 38^. Cells were tested negative for mycoplasma. Cells were mixed (ratio 1:1) and plated at 8×10^3^ per well in a μClear 96-well plate (Greiner Bio-One). The indicated SARS-CoV-2 strains were incubated with mAb, sera or nasal swabs at the indicated concentrations or dilutions for 15 minutes at room temperature and added to S-Fuse cells. The nasal swabs and sera were heat-inactivated 30 min at 56°C before use. 18 hours later, cells were fixed with 2% PFA, washed and stained with Hoechst (dilution 1:1,000, Invitrogen). Images were acquired with an Opera Phenix high content confocal microscope (PerkinElmer). The GFP area and the number of nuclei were quantified using the Harmony software (PerkinElmer). The percentage of neutralization was calculated using the number of syncytia as value with the following formula: 100 x (1 – (value with serum – value in “non-infected”)/(value in “no serum” – value in “non-infected”)). Neutralizing activity of each serum was expressed as the half maximal effective dilution (ED50). ED50 values (in μg/ml for mAbs and in dilution values for sera) were calculated with a reconstructed curve using the percentage of the neutralization at the different concentrations. We previously reported a correlation between neutralization titres obtained with the S-Fuse assay and a pseudovirus neutralization assay ^45^.

### Clinical history of the patient infected with AY.4.2

A nasopharyngeal swab collected from a 10-year-old boy tested positive for SARS CoV-2 on October 20th 27, was sent to Henri Mondor sequencing platform in the context of a nationwide survey (called flash). Briefly, private and public diagnostic laboratories in France participate weekly to the national SARS CoV-2 genomic surveillance by providing a random subsampling of positive SARS CoV-2 samples to national sequencing platforms ^46^.

### SARS CoV-2 Sequencing of the patient infected with AY.4.2

The full-length SARS-CoV-2 genome from the patient was sequenced by means of next-generation sequencing. Briefly, viral RNA was extracted from nasopharyngeal swabs in viral transport medium using. Sequencing was performed with the Illumina COVIDSeq Test (Illumina, San Diego, California), that uses 98-target multiplex amplifications along the full SARS-CoV-2 genome. The libraries were sequenced with NextSeq 500/550 High Output Kit v2.5 (75 Cycles) on a NextSeq 500 device (Illumina). The sequences were demultiplexed and assembled as full-length genomes by means of the DRAGEN COVIDSeq Test Pipeline on a local DRAGEN server (Illumina). AY4.2 was assigned using Pangolin before being submitted to the GISAID database ^47^.

### Virus strains

The variant strains were isolated from nasopharyngeal swabs on Vero cells and amplified by one or two passages on Vero cells. The delta strain was isolated from a nasopharyngeal swab of a hospitalized patient returning from India. The swab was provided and sequenced by the laboratory of Virology of Hopital Européen Georges Pompidou (Assistance Publique – Hopitaux de Paris). Both patients provided informed consent for the use of the biological materials. Titration of viral stocks was performed on Vero E6, with a limiting dilution technique allowing a calculation of TCID50, or on S-Fuse cells. Viruses were sequenced directly on nasal swabs, and after one or two passages on Vero cells. Sequences were deposited on GISAID immediately after their generation, with the following IDs: D614G: EPI_ISL_414631; B.1.1.7: EPI_ISL_735391; B.1.1.351: EPI_ISL_964916; B.1.617.2: ID: EPI_ISL_2029113.

### Flow Cytometry

For studies on infected cells, Vero cells were infected with the indicated viral strains at a multiplicity of infection (MOI) of 0.1. At 48h post-infection, cells were detached using PBS-EDTA and transferred into U-bottom 96-well plates (50,000 cell/well). For studies on transfected cells, HEK293T cells were transfected in suspension using lipofectamine 2000 as per manufacturer’s instruction (ThermoFischer), using 25% of phCMV-SARS-CoV2-spike and 75% of pQCXIP-Empty. 24h post-transfection, cells were detached using PBS-EDTA and transferred into U-bottom 96-well plates (50,000 cell/well). Cells were then incubated for 30 min at 4°C with the indicated mAbs (1 μg/mL) or Serum (1:300 dilution or as indicated for dose response) in MACS buffer (PBS, 5g/L BSA, 2mM EDTA). Cells were washed with PBS, and stained using anti-IgG AF647 (1:600 dilution in MACS, 30 min at 4°C) (ThermoFisher). Cells were then fixed using PFA 4%, 30 minutes. Data were acquired on an Attune Nxt instrument (Life Technologies).

For ACE2 binding, 293T cells transfected with S proteins for 24 hours as described above were stained with soluble biotinylated ACE2 diluted in MACS buffer at indicated concentrations (from 60 to 0.01 μg/ml) for 30 min at 4°C. The cells were then washed twice with PBS and then incubated with Alexa Fluor 647-conjugated streptavidin (Thermo Fisher Scientific, 1:400) for 30 min at 4°C. Cells were then fixed using PFA 4%, 30 minutes. Data were acquired on an Attune Nxt instrument (Life Technologies).

Analysis was performed with FlowJo 10.7.1 (Becton Dickinson).

### Antibodies

The four therapeutic antibodies were kindly provided by CHR Orleans. Human anti-SARS-CoV2 mAbs were cloned from S-specific blood memory B cells of Covid19 convalescents (Planchais et al, manuscript in preparation). Recombinant human IgG1 mAbs were produced by co-transfection of Freestyle 293-F suspension cells (Thermo Fisher Scientific) as previously described ^48^ and purified by affinity chromatography using protein G sepharose 4 fast flow beads (GE Healthcare).

### Statistical analysis

Flow cytometry data were analyzed with FlowJo v10 software (Becton Dickinson). Calculations were performed using Excel 365 (Microsoft). Figures were drawn on Prism 9 (GraphPad Software). Statistical analysis was conducted using GraphPad Prism 9. Statistical significance between different groups was calculated using the tests indicated in each figure legend.

## Results

### Antibody recognition of the AY.4.2 variant spike

To characterize the function of the AY.4.2 spike, we introduced the T95I, Y145H and A222V signature mutations in an expression plasmid coding for the Delta protein^30^. We first examined the ability of the Delta and AY.4.2 spikes to bind to a panel of 14 anti-SARS-CoV-2 monoclonal antibodies targeting either the RBD or the NTD. We tested 4 clinically approved antibodies, Bamlanivimab (LY-CoV555), Etesevimab (LY-CoV016), Casirivimab (REGN10933) and Imdevimab (REGN10987) targeting the RBD ^31,32^ as well as 4 other anti-RBD (RBD-48, RBD-85, RBD-98 and RBD-109) and 6 anti-NTD (NTD-18, NTD-20, NTD-32, NTD-45, NTD-69 and NTD-71) antibodies derived from convalescent individuals (Planchais et al, in preparation). Neutralizing anti-SARS-CoV-2 mAbs targeting the RBD can be classified into 4 main categories depending on their binding epitope ^33,34^. RBD-48 and RBD-85 belong to the first category (‘Class 1’) and act by blocking binding of the ‘up’ conformation of RBD to ACE2 ^34^. The precise epitopes of RBD-98 and RBD-109 are not yet defined but overlap with those of RBD-48 and RBD-85. Casirivimab and Imdevimab are mixed in the REGN-COV2 cocktail from Regeneron (Ronapreve^™^) and target different domains of the RBD. Casirivimab is a Class 1 antibody whereas Imdevimab binds to a lateral domain and belongs to the Class 3 ^32^. The anti-NTD antibodies bind uncharacterized epitopes within this domain, as assessed by Elisa (not shown).

We previously assessed the ability of most of these antibodies to recognize the spikes of the Alpha, Beta and Delta variants ^3,30^. To study their activity against AY.4.2, we first transfected the plasmids expressing Delta and AY.4.2 into 293T cells and analyzed antibody binding by flow cytometry (Fig. 1a). In line with our previous results, the Delta variant was recognized by 9 of the 16 antibodies ^3,30^. AY.4.2 displayed the same binding profile as Delta (Fig. 1a).

**Figure 1.**
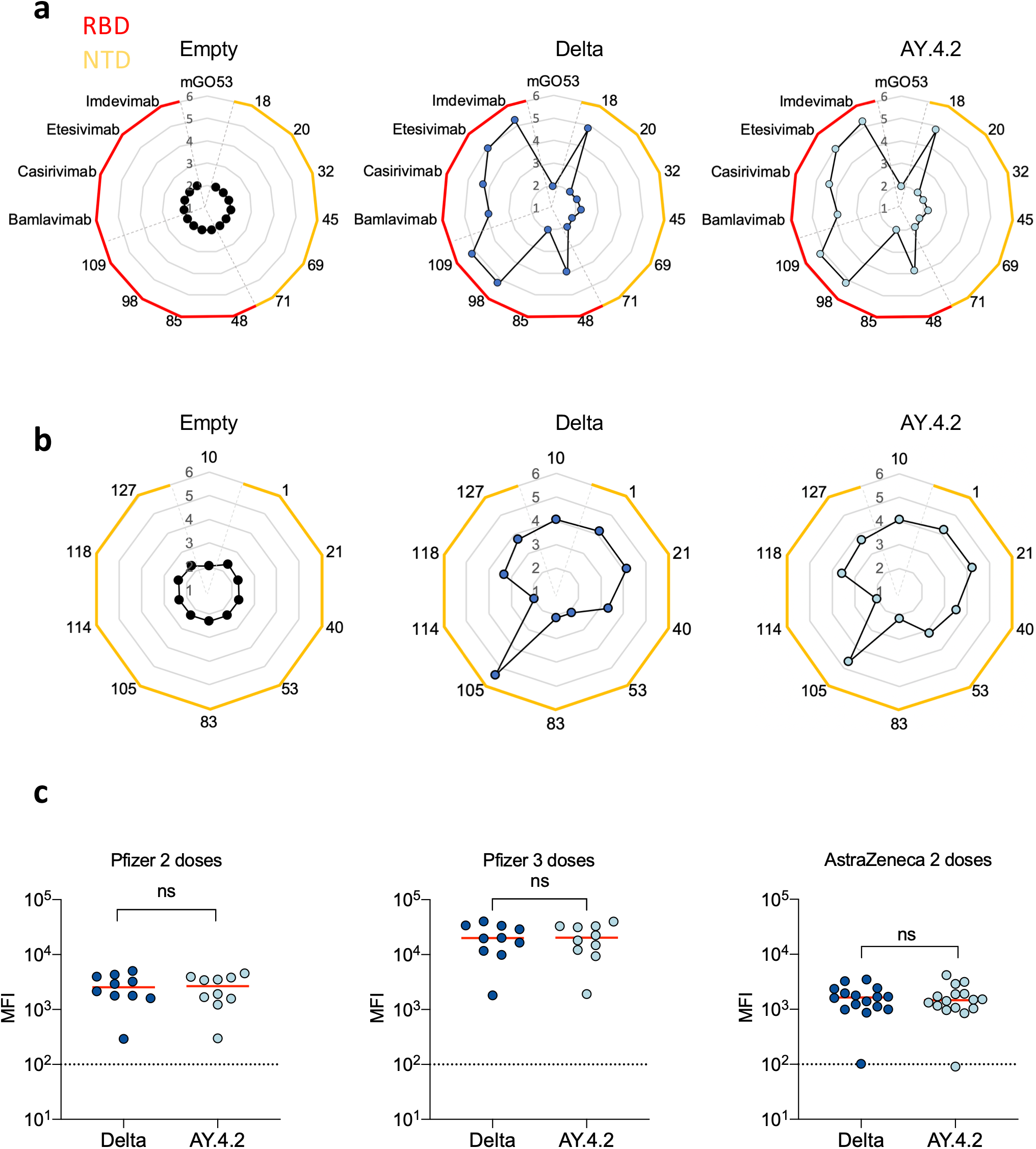
Antibody binding to cells expressing the Delta or AY.4.2 spike protein. (A-C) HEK293T cells were transiently transfected with plasmids expressing the Delta or AY.4.2 spike protein. (A-B) Binding of a panel of monoclonal antibodies targeting either the spike NTD or RBD. After 24h, cells were stained with the indicated antibody (1ug/mL). Radar charts represent for each antibody the logarithm of the median fluorescent intensity of the staining. Data are representative of two independent experiments (C) Binding of a panel of sera from vaccinated individuals. Sera from Pfizer vaccinated recipients were sampled at month 7 (M7) post- 2^nd^ dose (n=9) and at month 8 (M8), 1 month post-third dose (n=9). Sera from AstraZeneca vaccinated individuals were sampled at M5 post full vaccination. After 24h, cells were stained with Sera (1:300 dilution). A Wilconxon paired t-test was used to compare the two spike proteins.

Since the three mutations lie in the NTD, we extended our analysis to nine additional monoclonal antibodies targeting this domain. These antibodies were also cloned from SARS-CoV-2 infected individuals and bind to uncharacterized epitopes (Planchais et al, in preparation). As a control we used mAb10, a pan-coronavirus antibody that targets an unknown but conserved epitope within the S2 region ^21^ (Planchais, manuscript in preparation). They do not display any neutralizing activity against the ancestral Wuhan SARS-CoV-2 (not shown). Six out of the nine antibodies bound to the Delta and AY.4.2 spikes expressed at the cell surface, with various intensities (Fig. 1b). There was no major difference in their binding to Delta and AY.4.2 spikes, except for NTD-53 which bound slightly more to AY.4.2 than to Delta and, conversely, NTD-105 which bound slightly more to Delta than to AY.4.2 (Fig. 1b).

We next examined the binding of anti-spike antibodies present in the sera of vaccines to Delta and AY.4.2. We selected individuals that received either two doses of Pfizer vaccine, sampled 7 months post second dose (n=10), or three doses, sampled at least one month after the third dose (n = 9) (Table S1A). We also studied individuals immunized with two doses of AstraZeneca vaccine, sampled at 5 months post second dose (n=18) (Table S1B). Sera were tested at a 1:300 dilution, which allows a quantitative assessment of the antibody levels by flow cytometry ^35 36^. Overall antibody levels were similar after two doses of Pfizer or AstraZeneca vaccines, and increased by 8 fold after the boost of Pfizer vaccine (Fig. 1c). There was no major difference in the binding to the Delta and the AY.4.2 spikes (Fig. 1c). We then performed a titration of the antibody levels in a subset of 8 sera by serial dilutions and obtained similar binding titres for the two spikes (Fig. S1a), confirming the results obtained at the 1:300 dilution.

Altogether, these results indicate that the T95I, Y145H and A222V mutations are not associated with significant changes in recognition of the spike by a panel of 24 monoclonal antibodies and by sera from vaccine recipients.

### Fusogenicity and ACE2 binding of the AY.4.2 variant spike

We previously established a quantitative GFP-Split based cell-cell fusion assay to compare the fusogenic potential of mutant or variant spike proteins ^30,37^. In this assay, 293-T cells expressing part of the GFP protein (GFP1-10) are transfected with the spike plasmid. The transfected donor cells are then co-cultured with acceptor Vero cells expressing the other part of GFP (GFP11) ^30^. Upon cell-cell fusion, the syncytia become GFP+ and the fluorescent signal is scored with an automated confocal microscope ^30,37^. Of note, 293T cells were chosen as donors because they lack ACE2 and do not fuse with each other upon spike expression. Vero cells were selected as targets because they endogenously express ACE2 and are naturally sensitive to SARS-CoV-2. We thus analyzed the fusogenic activity of the AY.4.2 spike and compared it to D614G, Alpha and Delta variants. As previously reported, the D614G and Alpha spike variants were less fusogenic than Delta (Fig. 2a). With Delta, the area of syncytia was higher (Fig. 2a) despite similar levels of spike expressed at the surface of 293T donor cells (Fig. 2b, Fig S2a,b). The combination of T95I, Y145H and A222V substitutions did not modify the fusogenic activity of the Delta spike (Fig. 2a).

**Figure 2.**
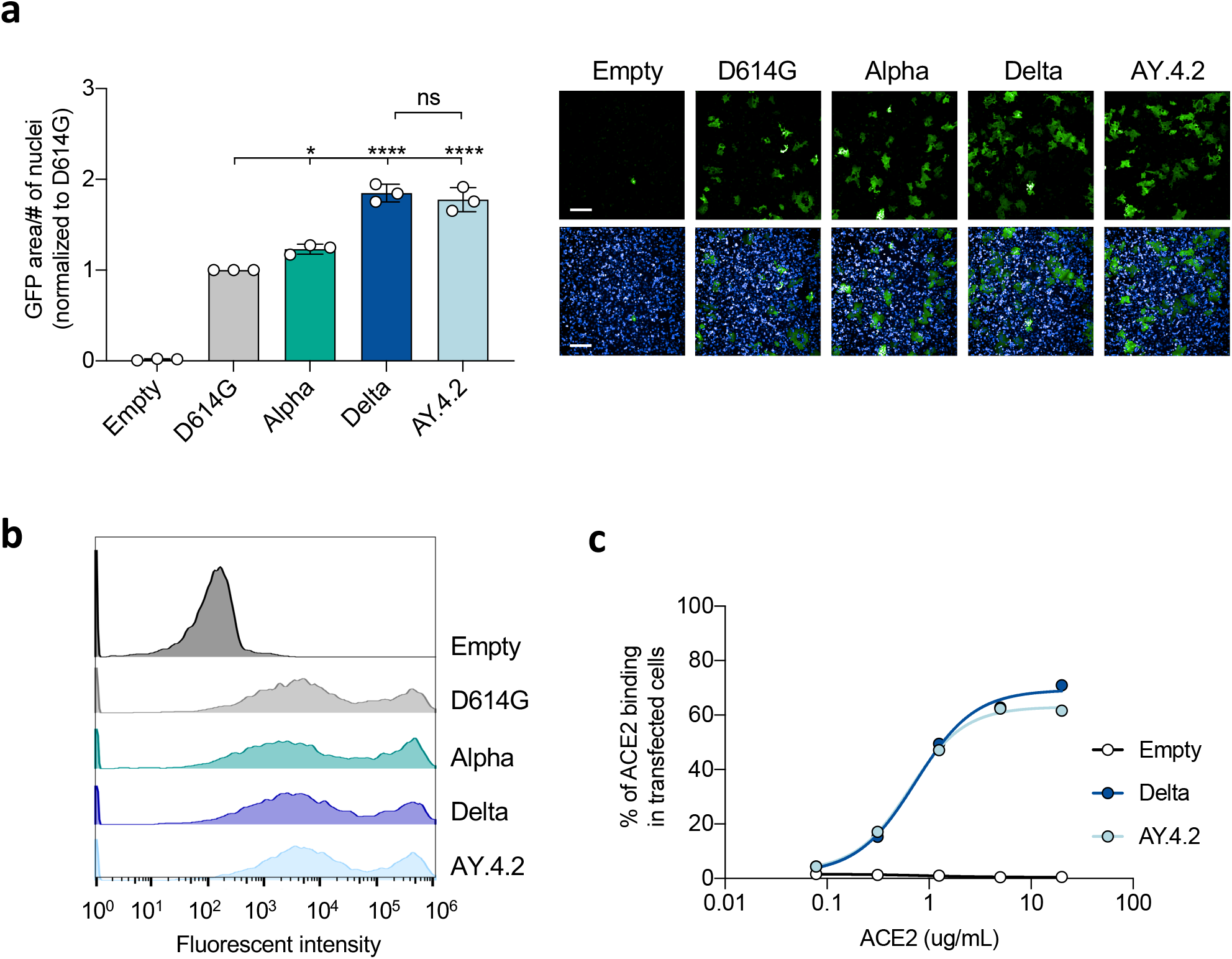
Comparison of Delta and AY.4.2 fusogenicity and ACE2 affinity. (A-B) Donor 293T GFP1-10 cells were transfected with the indicated spike encoding plasmid. (A) Donor cells were added to Vero GFP11 acceptor cells to assess fusion, using an Opera Phenix microscope (Perkin Elmer). Left Panel: Fusion was quantified by using the total GFP area/number of nuclei before normalizing to D614G for each experiment. Data are mean ± SD of three independent experiments. Statistical analysis: One-way ANOVA, each strain is compared to D614G or delta. ns: non-significant, ***P < 0.001. Right Panel: Representative images. Green: GFP-Split, Blue: Hoechst. Scale bars: 200 μm. (B) Donor cells were surface stained with a monoclonal anti-S antibody (129) to quantify spike expression. The data was then acquired by flow cytometry. Representative histogram of three independent experiments. (C) 293T cells were transfected with the indicated spike encoding proteins. After 24 h, they were stained with biotinylated ACE2 and fluorescent streptavidin before analysis by flow cytometry. Data are representative of two independent experiments.

We next explored AY.4.2 spike binding to the ACE2 receptor. To this aim, we transiently expressed the Delta and AY.4.2 proteins in 293T cells. Cells were then stained with a serial dilution of soluble biotinylated ACE2 and revealed with fluorescent streptavidin before analysis by flow cytometry (Fig. 2c). We previously reported using this assay that the spike protein of Alpha had the highest affinity to ACE2, followed by Delta and then by D614G, ^3,30^. Titration binding curves were generated with Delta and AY.4.2, showing no difference between the spikes’ affinity for ACE2 (Fig. 2c).

Therefore, the fusogenicity and ACE2 binding of the AY.4.2 spike are similar to the ones of the parental Delta variant.

### Isolation and characterization of an infectious AY.4.2 strain

We isolated the AY.4.2 variant from the nasopharyngeal swab of a symptomatic individual from the Paris region. The isolate was amplified by two passages on Vero E6 cells. Sequences of the swab and the outgrown viruses were identical and identified the AY.4.2 variant (GISAID accession ID: EPI_ISL_5748228, also termed hCoV-19/France/GES-HMN-21102260073/2021) (Fig. S3a). In particular, the spike protein contained the 3 expected mutations in the NTD (T95I, Y145H and A222V) when compared to the Delta strain used here as a reference. Viral stocks were titrated using S-Fuse reporter cells and Vero cells ^37,38^. S-Fuse cells allow rapid titration and measurement of neutralizing antibodies. They generate a GFP signal as soon as 6 hours post infection and the number of GFP+ cells correlates with the viral inoculum ^37 38^. Viral titres were similar in the two target cells and reached 10^5^ to 10^6^ infectious units/ml for the two strains. Syncytia were observed in infected Vero and S-Fuse cells (not shown). As expected, the syncytia were positive for spike staining (not shown).

We asked whether the spike present at the surface of infected cells displays the same characteristics as upon expression by transfection. We examined by flow cytometry the binding of neutralizing and non-neutralizing monoclonal antibodies to Vero cells infected with the Delta and AY.4.2 isolates. We observed the same profile of binding (Fig. 3a,b) for the two strains, and no noticeable difference with transfected 293T cells.

**Figure 3.**
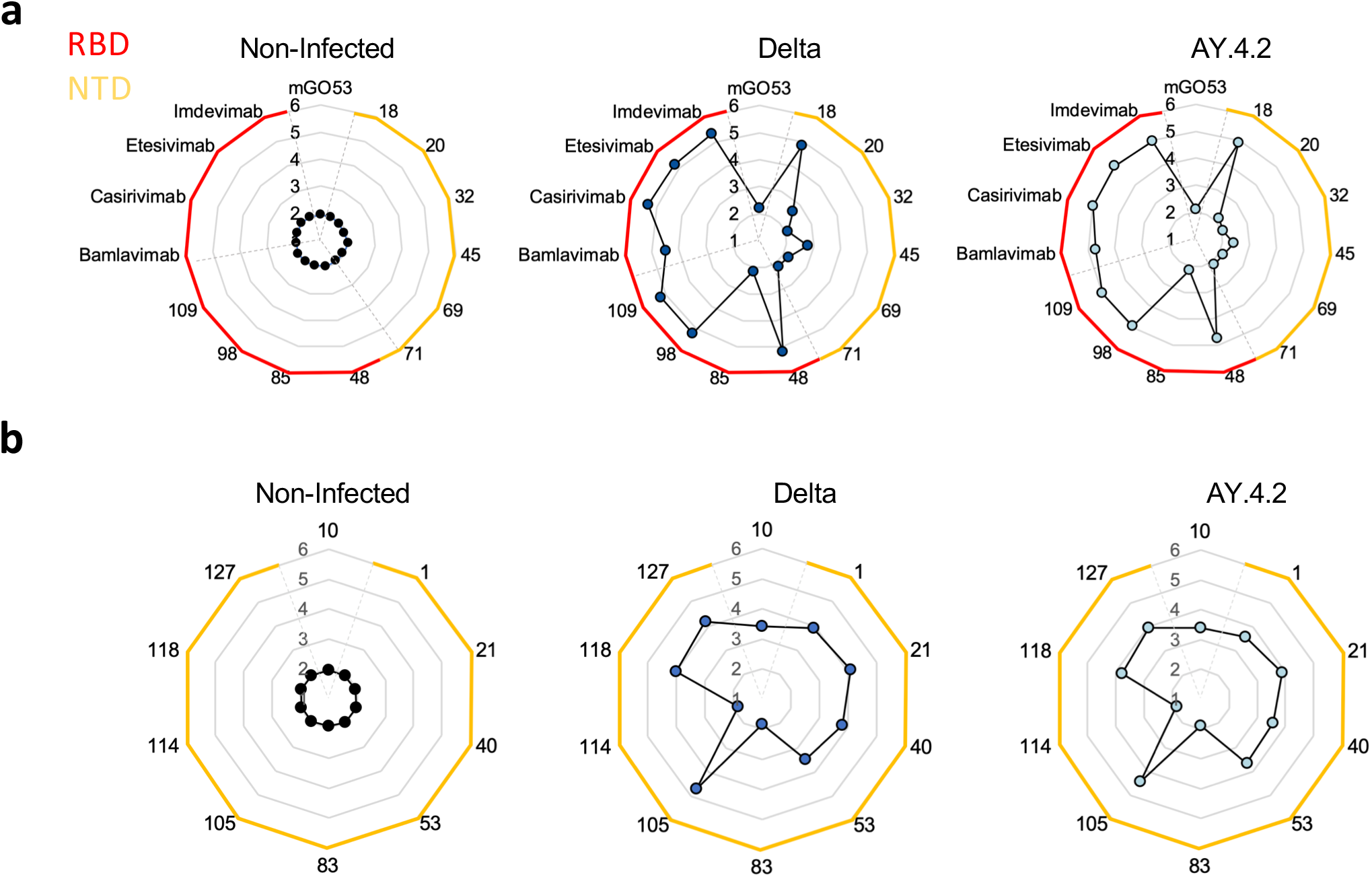
Antibody binding of the two viral isolated. (A-B) Vero cells were transiently infected with the two viral isolates, and harvested 48 hours post-infection for surface staining. (A-B) Binding of a panel of monoclonal antibodies targeting either the spike NTD or RBD, four of which are commercial therapeutic antibodies (1ug/mL). Radar charts represent for each antibody the logarithm of the median fluorescent intensity of the staining.

Altogether, these results indicate that the profile of antibody binding is similar in spike-expressing transfected 293-T cells and Vero infected cells. AY.4.2 and Delta infected cells display the same affinity to the panel of monoclonal antibodies we tested.

### Neutralization of AY.4.2 by monoclonal antibodies

We next compared the sensitivity of Delta and AY.4.2 strains to the previously described panel of neutralizing mAbs using the S-Fuse assay (Fig. 4a). 8 out of 14 antibodies neutralized both strains. With most of the neutralizing antibodies, we observed a slightly increased IC50s against AY.4.2 (average 2.2 fold increase when compared to Delta, Fig. 4a and Table S2). Bamlanivimab was inactive against AY.4.2, in agreement with previous results with Delta ^3 5 39^. Imdevimab displayed an incomplete neutralization. The maximum neutralization plateaued at 60% against AY.4.2, even at high antibody concentrations (1 μg/mL), whereas it reached almost 100 % against Delta (Fig. 4a). We obtained similar results with two different batches of Imdevimab (not shown). Therefore, AY.4.2 displays a slightly more elevated resistance to neutralization by the monoclonal antibodies tested than Delta. This resistance is more marked for Imdevimab.

**Figure 4.**
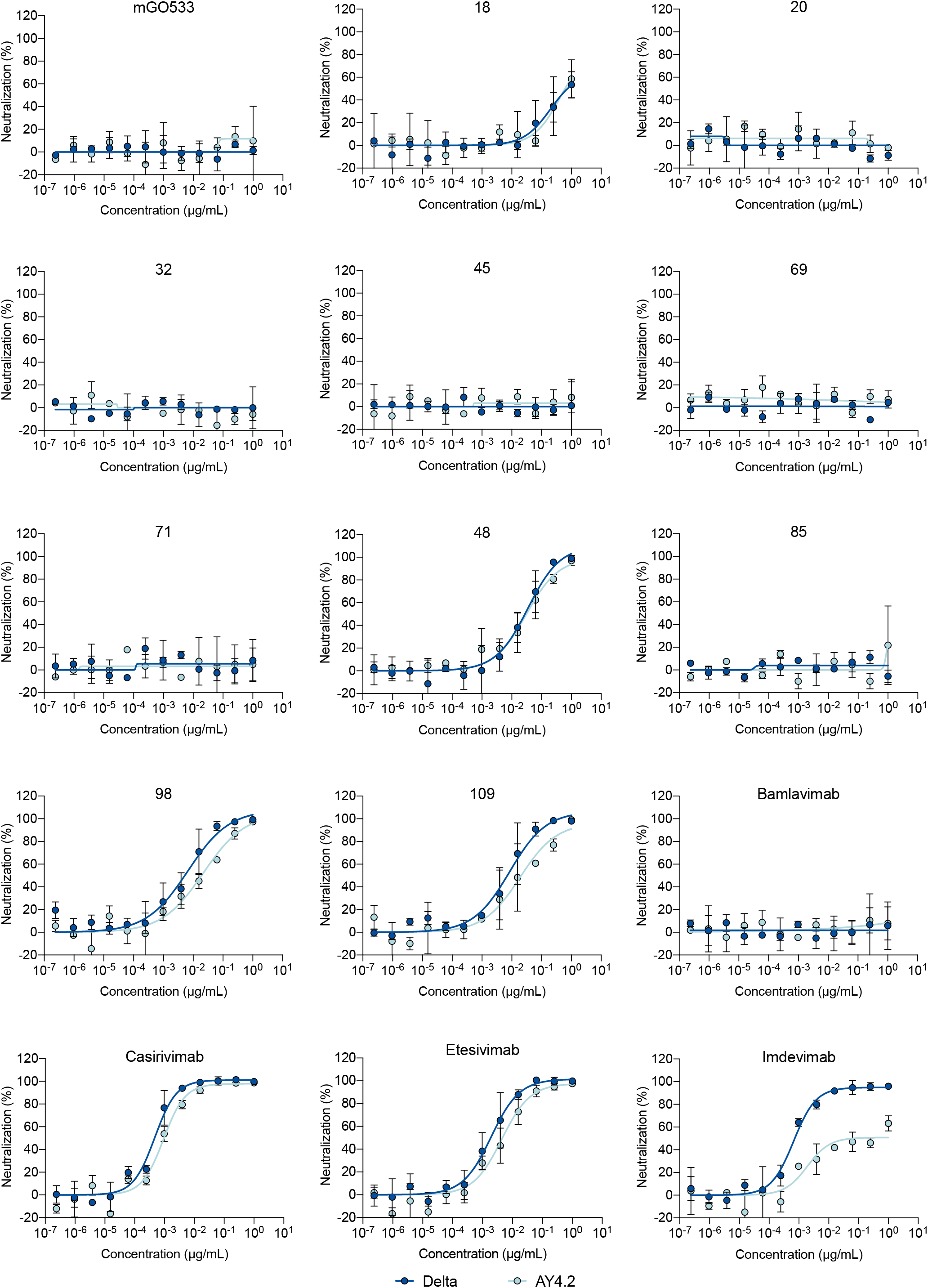
Neutralizing activity of monoclonal antibodies against the two viral isolates. (A) Dose response analysis of the neutralizing activity of a panel of monoclonal antibodies targeting either the spike NTD or RBD, four of which are commercial therapeutic antibodies. Data are mean ± SD of two independent experiments.

### Sensitivity of AY.4.2 to sera from vaccine recipients

We next asked whether vaccine-elicited antibodies neutralized AY.4.2. We used the same set of sera that were characterized by flow cytometry in Fig. 1 and compared their neutralizing activity against Delta and AY.4.2.

With the Pfizer vaccine, seven months after the second dose, the levels of neutralizing antibodies were relatively low against Delta (median ED50 of neutralization of 47), reflecting the waning of the humoral response at this time point ^3^ (Fig. 5a). These titres were lower against AY.4.2 (ED50 of 28, corresponding to a 1.7 fold decrease compared to Delta). One month after the booster dose (administrated at M7 post vaccination), titres strongly increased (25-50 fold), reaching 2716 and 1260 for Delta and AY.4.2 strains, respectively (Fig. 5a). We observed a 2.1 fold reduction in the neutralization titres against AY.4.2 when compared to Delta (Fig. 2b).

**Figure 5.**
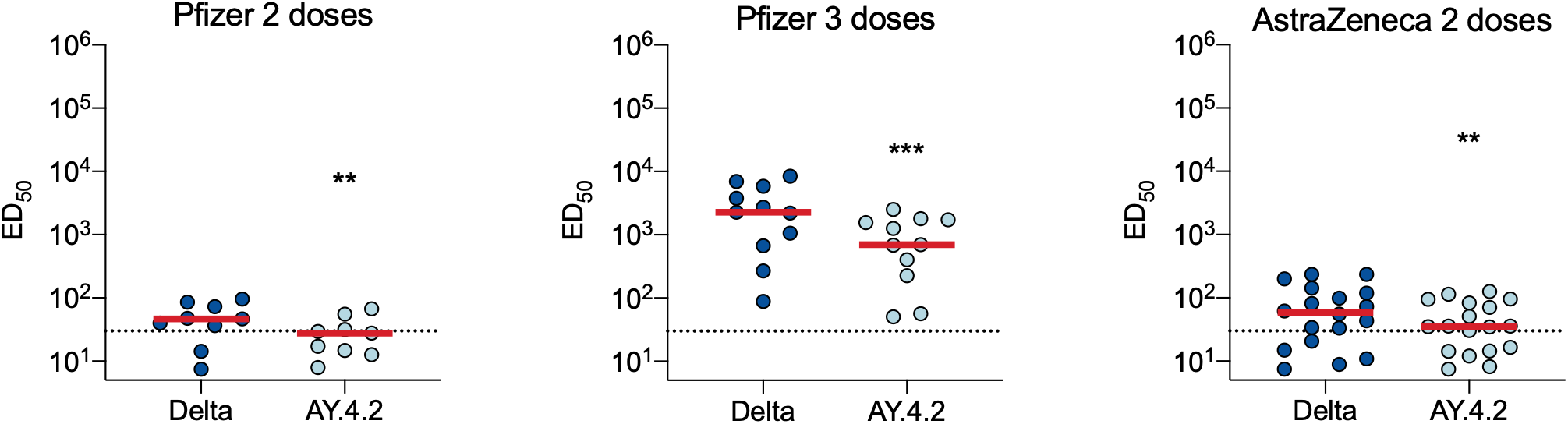
Neutralizing activity of vaccinated individuals’ sera against the two viral isolates. (A) ED50 of neutralization of the two viral isolates. Sera from Pfizer vaccinated recipients were sampled at month 7 (M7) post-2^nd^ dose (n=9) and at month 8 (M8), 1 month post-third dose (n=9). Sera from AstraZeneca vaccinated individuals were sampled at M5 post full vaccination. Data are mean from two independent experiments. A Wilconxon paired t-test was used to compare the two viral strains.

A similar pattern was observed with the AstraZeneca vaccine. Five months after the second dose, the neutralizing titres against Delta and AY.4.2 were low (ED50 of 58 and 35, respectively, representing a 1.6 decrease between Delta and AY.4.2).

Therefore, by using a set of sera with either low or high neutralizing antibody titres, we consistently observed a slight but significant decrease (1.7 to 2.1 fold reduction) of their activity against AY.4.2, when compared to the parental Delta variant.

## Discussion

Diversification of the Delta variant is regularly reported. The AY.4.2 sublineage was first identified in July 2021 and accounted for 15% and 20% sequenced Delta cases in UK, during the first and third weeks of November, respectively ^14^. This corresponds to an AY.4.2 logistic growth rate of 15% per week in this country ^14^. AY.4.2 has also been detected in dozens of countries. AY.4.2 was slowly but continuously rising and may thus display a slight selective advantage compared to the parental Delta strain. An increase of the growth rate may depend on the context and should not be necessary interpreted as a change in biological transmissibility ^14^. Preliminary lines of evidence indicated that hazard ratios for hospitalization or death are similar for Delta and AY.4.2, indicating that outcomes of AY.4.2 are not more severe than Delta cases ^14^. Since December 2021, Delta and its sublineages, including AY.4.2, has been replaced by Omicron. It remains however of interest to understand the parameters that may have favoured the spread of AY.4.2, compared to the original Delta strain.

The AY.4.2 strain remains poorly characterized. It carries 3 main substitutions, T95I, Y145H and A222V, compared to the parental Delta lineage. Here, we show that the AY.4.2 spike is functionally very close to that of Delta. By using a panel of 24 monoclonal antibodies targeting either the RBD or the NTD, we did not detect major differences in antibody recognition, when the spikes are expressed by transient transfection in 293-T cells. Polyclonal sera from individuals having received either Pfizer or AstraZeneca vaccines similarly recognized the two spikes. Their fusogenic activity, when measured in a syncytia formation assay ^30^ and the binding affinity to ACE2 were also similar for AY.4.2 and Delta.

We isolated an authentic AY.4.2 strain from an infected patient and examined its sensitivity to antibody neutralization. Future work in more relevant models, such as primary human bronchial epithelium ^40^ or viral competition experiments will help determining whether AY.4.2 is more fit than the parental lineage in culture systems. We analyzed the profile of binding of a panel of monoclonal antibodies to infected cells. We did not observe major differences between the two strains.

We then studied the neutralization of the two viral isolates by a panel of monoclonal antibodies. Imdevimab, a therapeutic antibody used in combination with Casirivimab in the commercially approved REGN-COV2 cocktail from Regeneron and Roche (Ronapreve^™^), incompletely neutralized AY.4.2. Even at high concentration, the neutralizing activity plateaued at 60%. Incomplete neutralization and deviation from sigmoidal neutralization curves have been for instance previously observed with some HIV broadly neutralizing antibodies (bNAbs) ^41^. This process has been attributed to heterogeneity in glycosylation of the HIV gp120/gp41 Env complex ^41^. Our results strongly suggest that the various conformations or glycosylation of the AY.4.2 spike at the virion surface may display different sensitivities to antibody neutralization.

As AY4.2 does not harbour mutations within the epitope of Imdevimab, our results also indicate that mutations in the NTD of the spike may remotely impact the accessibility of anti-RBD antibodies. The 3D structure of the spike shows that some regions of the NTD are in close proximity to the RBD ^34 42^. Imdevimab binds to a lateral region of the RBD and (Class 3 antibody) and may thus be more affected by changes in the NTD than other anti-RBD antibodies binding to the apex of the spike. Furthermore, the other neutralizing anti-RBD antibodies that we tested displayed a slight decrease in sensitivity to AY4.2, when compared to Delta (1.8 to 3.3 fold increase of the IC50), except for antibody 48. Of the 6 monoclonal antibodies targeting the NTD we tested, only NTD-18^3^ neutralized Delta. NTD-18similarly neutralized both strains at high concentration. It will be worth determining whether other antibodies targeting the NTD super antigenic site ^32,43^ are less active against AY.4.2.

We further show that sera from individuals having received two or three doses of Pfizer vaccine, or two doses of AstraZeneca, remained active against AY.4.2, with however a 1.7 to 2.1 fold reduction in neutralization titres. This decrease may be attributed to the slight reduction of the efficacy of some Imdevimab-like antibodies in the serum, or targeting other RBD and NTD regions in the spike.

Preliminary epidemiology results of vaccine effectiveness in UK, for both symptomatic and non-symptomatic breakthrough infections, indicated no significant differences between AY.4.2 and non-AY.4.2 cases ^44^. Our results indicate that the slight decrease in neutralizing titres reported here did not significantly impact vaccine effectiveness against AY.4.2, at least within 6-7 months post-vaccination.

## Contributors

Experimental strategy design, experiments: NS, DP, WB, JB, CP, MP, IS, FGB, FP, AB, ESL, TB, OS Vital materials: CR, SF, CP, DV, HP, NR, MS, JP, MP, AS, LH, TP, HM, JP

Manuscript writing: NS, OS

Manuscript editing: NS, DP, CR, SF, TB, JP, OS

## Declaration of Interests

C.P., H.M., O.S, T.B. have a pending patent application for an anti-RBD mAb not used in the present study (PCT/FR2021/070522).

## Acknowledgments

We thank patients who participated to this study, members of the Virus and Immunity Unit for discussions and help, Nathalie Aulner and the UtechS Photonic BioImaging (UPBI) core facility (Institut Pasteur), a member of the France BioImaging network, for image acquisition and analysis. We thank Julien Puech, Julien Rodary et Dhiaeddine Edriss from the Hopital Européen Georges Pompidou for their help with sequencing. We thank Fabienne Peira, Vanessa Legros and Laura Courtellemont for their help with the cohorts.

Work in OS lab is funded by Institut Pasteur, Fondation pour la Recherche Médicale (FRM), Urgence COVID-19 Fundraising Campaign of Institut Pasteur, ANRS, the Vaccine Research Institute (ANR-10-LABX-77), Labex IBEID (ANR-10-LABX-62-IBEID), ANR/FRM Flash Covid PROTEO-SARS-CoV-2 and IDISCOVR. Work in UPBI is funded by grant ANR-10-INSB-04-01 and Région Ile-de-France program DIM1-Health. NS is supported by the French Ministry of Higher Education, Research and Innovation. DP is supported by the Vaccine Research Institute. HM lab is funded by the Institut Pasteur, the Milieu Intérieur Program (ANR-10-LABX-69-01), the INSERM, REACTing, EU (RECOVER) and Fondation de France (#00106077) grants. ESL lab is funded by Institut Pasteur and the French Government’s Investissement d’Avenir programme, Laboratoire d’Excellence “Integrative Biology of Emerging Infectious Diseases” (grant n°ANR-10-LABX-62-IBEID). The funders of this study had no role in study design, data collection, analysis and interpretation, or writing of the article.

## Data Sharing Statement

All data supporting the findings of this study are available within the paper and are available from the corresponding author upon request.

## Figure Legends

**Table S1. Clinical Data regarding the Pfizer and Astra Zeneca vaccinated individuals.**

**Table S2. EC50 of neutralizing monoclonal antibodies**.

**Table S3. Primers used for Site Directed Mutagenesis and sequencing**.

**Figure S1. Antibody binding to cells expressing the Delta or AY.4.2 spike protein**.

(A) HEK293T cells were transiently transfected with plasmids expressing the different spike proteins. Dose response binding of a panel of sera from vaccinated individuals in the Astra Zeneca cohort.

**Figure S2. Comparison of Delta and AY.4.2 fusogenicity and ACE2 affinity**.

(A-B) Donor 293T GFP1-10 cells were transfected with the indicated spike encoding plasmid. (A) Donor cells were surface stained with a monoclonal anti-S antibody (129) to quantify spike expression. The data was then acquired by flow cytometry. Left panel: Percentage of positive cells. Right panel: Median fluorescent intensity in the positive cells. Data are mean ± SD of three independent experiments. Statistical analysis: One-way ANOVA, each strain is compared to D614G or delta. ns: non-significant

**Figure S3. Comparison of the Delta and the AY.4.2 strains**.

(A) Schematic overview of the Delta and AY4.2 sublineages consensus sequences with a focus on the spike built with the Sierra tool 58. Amino acid modifications in comparison to the ancestral Wuhan-Hu-1 sequence (NC_045512) are indicated.

## Notes

### Competing Interest Statement

C.P., H.M., O.S, T.B. have a pending patent application for an anti-RBD mAbs not used in the present study (PCT/FR2021/070522).

