## Supplementary Data for "Fusogenicity and neutralization sensitivity of the SARS-CoV-2 Delta sublineage AY.4.2"

### Supplementary Information

Table S1. Clinical Data regarding the Pfizer and Astra Zeneca vaccinated individuals……………………….. 2

Table S2. EC50 of neutralizing monoclonal antibodies……………………………………………………... 3

Table S3. Primers used for Site Directed Mutagenesis and sequencing. …………………………………… 4

Figure S1. Antibody binding to cells expressing the Delta or AY.4.2 spike protein. (A) HEK293T cells were transiently transfected with plasmids expressing the different spike proteins. Dose response binding of a panel of sera from vaccinated individuals in the Astra Zeneca cohort……………………………………… 5

Figure S2. Comparison of Delta and AY.4.2 fusogenicity and ACE2 affinity.(A-B) Donor 293T GFP1-10 cells were transfected with the indicated spike encoding plasmid. (A) Donor cells were surface stained with a monoclonal anti-S antibody (129) to quantify spike expression. The data was then acquired by flow cytometry. Left panel: Percentage of positive cells. Right panel: Median fluorescent intensity in the positive cells. Data are mean ± SD of three independent experiments. Statistical analysis: One-way ANOVA, each strain is compared to D614G or delta ns: non-significant…………………………………………………… 6

Figure S3. Comparison of the Delta and the AY.4.2 strains.(A) Schematic overview of the Delta and AY4.2 sublineages consensus sequences with a focus on the spike built with the Sierra tool 58. Amino acid modifications in comparison to the ancestral Wuhan-Hu-1 sequence (NC_045512) are indicated…………… 7

**a**

| ID | Sex | Age | Vaccine | 1st dose | 2nd dose | 3rd dose | M7 sampling post-2nd dose | | M1 sampling post-3rd dose | |
|---|---|---|---|---|---|---|---|---|---|---|
| | | | | | | | Date | Days | Date | Days |
| VAC PF #1 | Female | 64 | Pfizer | 7/Jan/21 | 28/Jan/21 | 1/Jul/21 | / | / | 31/Aug/21 | 61 |
| VAC PF #2 | Male | 69 | Pfizer | 7/Jan/21 | 28/Jan/21 | 31/Aug/21 | 31/Aug/21 | 215 | 28/Sep/21 | 28 |
| VAC PF #3 | Male | 60 | Pfizer | 8/Jan/21 | 29/Jan/21 | 31/Aug/21 | 31/Aug/21 | 214 | 12/Oct/21 | 42 |
| VAC PF #4 | Male | 65 | Pfizer | 6/Jan/21 | 28/Jan/21 | 6/Sep/21 | 26/Aug/21 | 210 | 8/Oct/21 | 32 |
| VAC PF #5 | Male | 53 | Pfizer | 6/Jan/21 | 26/Jan/21 | 31/Aug/21 | 30/Aug/21 | 216 | 6/Oct/21 | 36 |
| VAC PF #6 | Female | 36 | Pfizer | 6/Jan/21 | 28/Jan/21 | 16/Nov/21 | 31/Aug/21 | 215 | / | / |
| VAC PF #7 | Male | 74 | Pfizer | 6/Jan/21 | 29/Jan/21 | 13/Sep/21 | 7/Sep/21 | 221 | 11/Oct/21 | 28 |
| VAC PF #8 | Male | 59 | Pfizer | 6/Jan/21 | 27/Jan/21 | 31/Aug/21 | 31/Aug/21 | 216 | 6/Oct/21 | 36 |
| VAC PF #9 | Male | 52 | Pfizer | 7/Jan/21 | 28/Jan/21 | 3/Nov/21 | 31/Aug/21 | 215 | / | / |
| VAC PF #10 | Male | 62 | Pfizer | 8/Jan/21 | 29/Jan/21 | 31/Aug/21 | 31/Aug/21 | 214 | 5/Oct/21 | 35 |
| VAC PF #11 | Female | 63 | Pfizer | 31/Jan/21 | 31/Jan/21 | 6/Sep/21 | 3/Sep/21 | 215 | 11/Oct/21 | 35 |

**b**

| ID | Sex | Age | Vaccine | 1st dose | 2nd dose | M6 sampling post-2nd dose | |
|---|---|---|---|---|---|---|---|
| | | | | | | Date | Days |
| VAC AZ #1 | Female | 57 | AstraZeneca | 6/Mar/21 | 5/May/21 | 4/Oct/21 | 152 |
| VAC AZ #2 | Female | 64 | AstraZeneca | 11/Feb/21 | 5/May/21 | 1/Oct/21 | 149 |
| VAC AZ #3 | Male | 58 | AstraZeneca | 11/Feb/21 | 6/May/21 | 1/Oct/21 | 148 |
| VAC AZ #4 | Female | 55 | AstraZeneca | 11/Feb/21 | 6/May/21 | 30/Sep/21 | 147 |
| VAC AZ #5 | Female | 61 | AstraZeneca | 9/Feb/21 | 15/Apr/21 | 11/Oct/21 | 179 |
| VAC AZ #6 | Female | 63 | AstraZeneca | 15/Feb/21 | 5/May/21 | 5/Oct/21 | 153 |
| VAC AZ #7 | Female | 59 | AstraZeneca | 9/Feb/21 | 4/May/21 | 5/Oct/21 | 154 |
| VAC AZ #8 | Female | 59 | AstraZeneca | 12/Feb/21 | 19/May/21 | 8/Oct/21 | 142 |
| VAC AZ #9 | Male | 61 | AstraZeneca | 9/Feb/21 | 4/May/21 | 4/Oct/21 | 153 |
| VAC AZ #10 | Female | 61 | AstraZeneca | 10/Feb/21 | 5/May/21 | 1/Oct/21 | 149 |
| VAC AZ #11 | Female | 55 | AstraZeneca | 9/Feb/21 | 4/May/21 | 1/Oct/21 | 150 |
| VAC AZ #12 | Male | 73 | AstraZeneca | 26/Mar/21 | 28/May/21 | 8/Oct/21 | 133 |
| VAC AZ #13 | Male | 57 | AstraZeneca | 5/Feb/21 | 3/May/21 | 1/Oct/21 | 151 |
| VAC AZ #14 | Female | 60 | AstraZeneca | 14/Feb/21 | 4/May/21 | 30/Sep/21 | 149 |
| VAC AZ #15 | Female | 60 | AstraZeneca | 7/Apr/21 | 4/May/21 | 7/Oct/21 | 156 |
| VAC AZ #16 | Female | 63 | AstraZeneca | 12/Feb/21 | 7/May/21 | 7/Oct/21 | 153 |
| VAC AZ #17 | Male | 56 | AstraZeneca | 18/Feb/21 | 11/May/21 | 8/Oct/21 | 150 |
| VAC AZ #18 | Male | 64 | AstraZeneca | 7/Feb/21 | 19/May/21 | 6/Sep/21 | 110 |

**Table S1. Clinical Data regarding the (a) Pfizer and (b) Astra Zeneca vaccinated individuals.**

| Antibody | Delta EC50 | AY.4.2 EC50 | Fold change |
|---|---|---|---|
| 48 | 34.8 | 30.5 | 0.9 |
| 98 | 6.6 | 22.0 | 3.3 |
| 102 | 1.3 | 2.4 | 1.9 |
| 109 | 8.4 | 18.7 | 2.2 |
| Casirivimab | 0.5 | 0.9 | 2.0 |
| Etesivimab | 1.9 | 4.4 | 2.3 |
| Imdevimab | 0.7 | 1.9 | 2.9 |

**Table S2. EC50 of neutralizing monoclonal antibodies.**

**a**

| | |
|---|---|
| T95I_F | TTTGCCAGCATCGAGAAGTCC |
| T95I_R | GTACACGCCATCGTTGAAG |
| Y145H_F | GGACGTCTACCACCACAAGAACAAC |
| Y145H_R | AGGAAGGGGTCGTTGCAG |
| A222V_F | GGGATTCAGTGTGCTGGAACCCCTGGTG |
| A222V_R | TGTGGCAGATCGCGCACG |
| D614G_F | CTGTACCAGGGCGTGAATTGCACAGAGGTG |
| D614G_R | AACGGCCACCTGGTTGCT |

**b**

| | |
|---|---|
| phCMV For | CTCTTTCCTACAGCTCCTGG |
| phCMV Int1 For | AGCGAGTTCCGCGTGTACAG |
| phCMV Int1_Delta For | GCCACCAACGTGGTCATCAA |
| phCMV Int2 For | CGCAAGCGCATTAGCAACTG |
| phCMV Int3 For | GCGTGCTGACCGAGAGTAAC |
| phCMV Int4 For | GCAACCTGCTGCTGCAGTAC |
| phCMV Int5 For | GTGGTCAACCAGAACGCTCAG |
| phCMV Int6 For | AGAACCACACAAGCCCCGAC |
| phCMV Rev | TAGCCAGAAGTCAGATGCTC |

**Table S3. Primers used for (a) Site Directed mutagenesis (b) Sequencing**

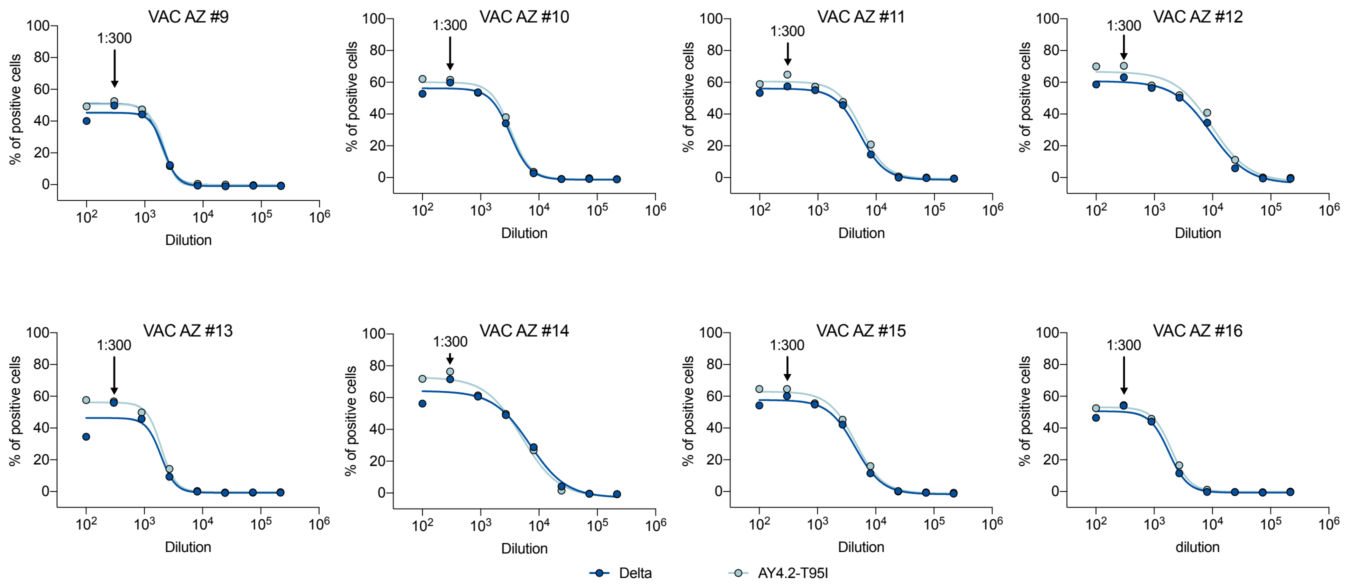


**Fig. S1. Dose response binding of sera from vaccinated individuals.**

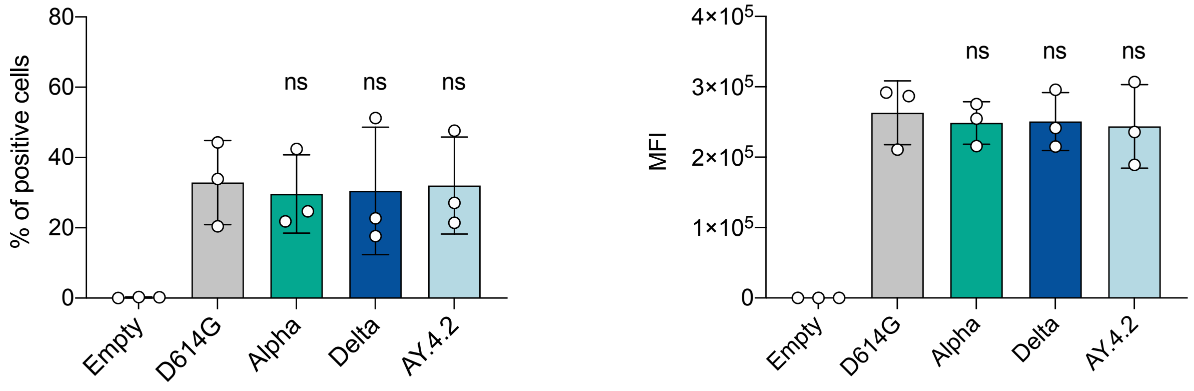


**Fig. S2. Comparison of Delta and AY.4.2 spike surface expression.**

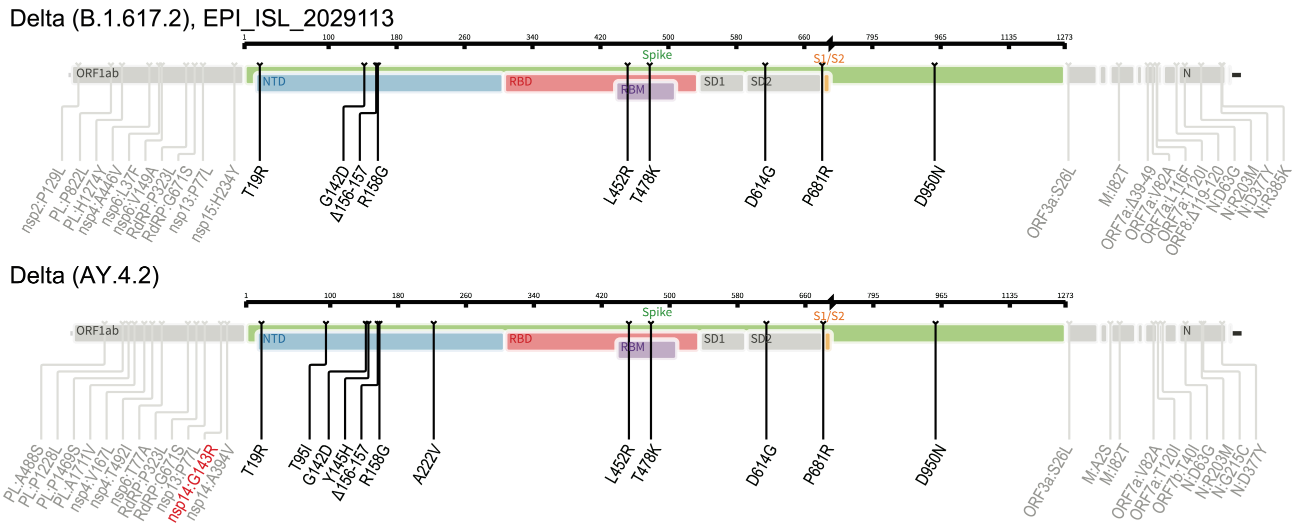


**Fig. S3. Comparison of the Delta and the AY.4.2 strains.**
